# Rapid adjustment of pecking trajectory to prism-induced visual shifts in crows

**DOI:** 10.1101/298414

**Authors:** Hiroshi Matsui, Ei-Ichi Izawa

## Abstract

**Summary statement:** The mechanisms underlying birds’ pecking skills are not known. We examined whether pigeons and crows adjust their pecking to the visual distortion caused by prisms. We found that vision plays a role only before movement begins in pigeons, whereas crows may have more flexible visuomotor skills.

**ABSTRACT:** Pecking in birds is analogous to reaching and grasping movements in primates. Although pecking in pigeons is highly stereotypic, crows show dexterous pecking skills. To unveil what sensorimotor mechanisms underlie the flexibility of pecking in crows, the current study examined whether pigeons and crows adjust their pecking to the visual distortion induced by prisms. Because prisms induce visual shifts of object positions, birds were required to adjust their movements. Pecking kinematics were examined before and after attaching prisms in front of the birds’ eyes. As a result, crows showed faster adjustment of pecking trajectories than pigeons. Correlational analysis showed that the initial deviation of trajectory strongly influenced the subsequent deviation in pigeons, but not in crows. These results suggest that pecking in pigeons is controlled in a feedforward manner, and vision plays a role before movement initiation, although pecking in crows is under the on-line motor control, serving a possible mechanism for flexible visuomotor skills.

## INTRODUCTION

Pecking behaviour is the fundamental motor repertoire in avian foraging, which is analogous to reaching and grasping movement in primates, and consists of two motor components, including head-reaching and bill-grasping (Bermejo et al., 1989; Delius, 1985). Despite the superficial similarities of these motor actions to primates’ arm-reaching and hand-grasping (Delius, 1985; Klein et al., 1985), it is still unknown whether avian pecking is controlled by similar visuomotor mechanisms, because of the clear anatomical differences between primates and birds. In primates, hands and arms (i.e. effector organs) are anatomically separated from the eyes on the head. This body structure enables primate eyes to see the locations of a target and their hands in a stable view, and to control the reaching movement using on-line visual feedback (Keele, 1968; Sarlegna and Mutha, 2015). However, in birds, both the bill and eyes are mounted together on the head, which causes an unstable and moving view associated with the head-reaching movement. This avian anatomy raises the question, what visuomotor mechanism controls avian pecking? Specifically, to what extent do birds use vision as a source of information to achieve accurate movements?

Previous studies on avian pecking have used pigeons (*Columba livia*) as subjects (Bout and Zeigler, 1984; Delius, 1985; Klein et al., 1985, LaMon and Zeigler, 1984; Theunissen et al., 2017; Zweers, 1982), and revealed the capability of visual control in their pecking. Pecking of pigeons consisted of three phases of movements: a fast standstill of head movement in front of a target food item (which is called ‘fixation’), initiation of head-reaching to the target, and grasping of the target by the bill (Delius, 1985). Once head-reaching has been initiated, pigeons typically close their eyes during pecking, suggesting that vision plays a role only in planning the reaching and grasping movements based on the target location and size determined during fixation, and vision is not involved after initiation of the movement (LaMon and Zeigler, 1984). Thus, avian pecking is possibly controlled in a feedforward manner, according to movements pre-planned during fixation, and performed without movement correction (Goodale, 1982; Theunissen et al., 2017). This ballistic mechanism to control pecking may be relevant to feeding ecology for pigeons to consume non-moving foods such as seeds on the ground. Pecking with feedforward motor control may be sufficiently effective, because it may not be necessary to peck at stable objects with visually-guided control during movement (henceforth, we refer this kind of motor control as ‘the on-line motor control’).

In contrast, a recent study suggested the involvement of on-line motor control in pecking of crows, omnivorous/carnivorous species (Matsui and Izawa, 2017). Matsui and Izawa (2017) compared motor learning between large-billed crows (*Corvus macrorhynchos*) and pigeons by attaching artificial extensions to their bills. Because the bill extensions distorted the timing of contact to a food, birds were required to adjust their grasping timing to successfully grasp food items. The experiment yielded distinct results between these two species: crows rapidly adapted to the bill extensions within a few trials, whereas pigeons did not succeed in obtaining food items with the extended bills. The rapid learning of successful pecking with the bill extension suggested pecking of crows might be under on-line control based on vision. However, there has been no behavioural evidence of the involvement of vision during pecking in birds such as crows.

The purpose of the current study was to investigate the involvement of vision during pecking in crows in comparison to pigeons. The involvement of vision was examined by using prism-induced visual shifts. Because prism glasses laterally shifted visual images on the retina, subjects were required to adjust their movement in the opposite direction of the visual distortion. Based on the previous studies described above, we hypothesized that pigeons would use the feedforward mechanism in their pecking, and in this mechanism the role of vision is limited before movement initiation. In contrast, under the on-line control mechanism, which was suggested in crows, vision plays a role during movements. Based on these motor control mechanisms, we predicted that we would see distinct results in responses to prism-induced visual shifts in pigeons and crows. In pigeons, we predicted that the feedforward mechanism would result in gradual adjustment to the prism-induced visual shifts, because sensory teaching signals would be obtained after movements were completed. However, if crows have the on-line control mechanism, we predicted that learning would be rapidly completed, because on-line control enables organisms to achieve deviations of movement trajectories induced by prisms. In addition, we predicted that the pecking trajectories of crows and pigeons would be very different, based on these motor control mechanisms. Under feedforward motor control, deviation of a trajectory at a given time point would influence subsequent deviation, because the trajectory is not corrected during movements. Thus, we predicted that the correlation between trajectory deviations would be high under feedforward motor control. We predicted that this would not be the case for on-line motor control, because the deviation of the trajectory could be corrected during movements. Therefore, we predicted that under on-line motor control (i.e. in crows), there would be less correlation between trajectory deviations than under feedforward motor control (i.e. in pigeons).

## MATERIALS AND METHODS

### Subjects and housing

Three adult pigeons (unknown sex, body weight: 290 – 349 g) and three sub-adult large-billed crows (three females, body weight: 510 – 715 g) were used. All birds were experimentally naive and wild-caught in Tokyo, as authorized by the Environmental Bureau of Tokyo Metropolitan Government (Permission #4005). Pigeons and crows were kept in different rooms, and housed individually in stainless steel-mesh home cages (*W* 35 cm × *D* 30 cm × *H* 35 cm for pigeons, *W* 43 cm × *D* 60 cm × *H* 50 cm for crows) for approximately one month for the experimental period, plus three days for acclimation to the experimental chamber. Conspecific individuals were placed side-by-side to allow them visual and audio-vocal social communication with one another. During the experimental period, crows were regularly transferred into an outdoor aviary (*W* 1.5 m × *D* 2.8 m × *H* 1.7 m) for 2 – 3 hours after daily experimental sessions, to allow crows to bathe and have direct social interactions with other conspecifics, although crows were not provided with food in the outdoor aviary. After all the experiments were finished, birds were transferred back to relatively large outdoor aviaries for group housing (*W* 3 m^2^ × *H* 1.5 m for pigeons, 100 m^2^ × *H* 3 m for crows), and used for other behavioural studies. Mixed grains with mineral supplements were fed to pigeons as daily diets, and dry foods, cheese, and eggs were fed to crows, respectively. During the experimental period, no food was provided to the pigeons and crows for five hours before the daily experimental session, and sufficient food was provided after the session. Water was freely available in the home cages. The room was maintained at 21 ± 2°C in a 13 L:11 D cycle, with light onset at 08:00. The experimental and housing protocols adhered to Japanese National Regulations for Animal Welfare, and were approved by the Animal Care and Use Committee of Keio University (no. 14077).

### Apparatus

The experiment was conducted in an experimental chamber (*W* 35 cm *× D* 30 cm *× H* 35 cm for pigeons, *W* 68.5 cm *× D* 62 cm *× H* 180.5 cm for crows; Figure 1). The chamber for crows consisted of a platform table (*W* 39 cm × *D* 19 cm and 10 cm above the floor). Given the different behavioural patterns of pecking under natural feeding situations, food used as pecking targets was presented on the table in different ways to pigeons and crows in each trial. For pigeons, which feed seeds and grains typically by sequential pecking, an array of five corns was presented at five specific positions in a line with 5 cm intervals on a frontal wall, 13 cm above the floor of the chamber (Figure 1a). The foods were attached with moderately adhesive tapes, from which pigeons could pick up the foods. For crows, which typically feed by a non-sequential single-shot pecking at a target, a small piece of cheese (approx. less than 1 cm sphere) was attached to the tip of a metal wire to lift it 5 cm above the platform table (Figure 1b). A high-speed camcorder (300 frame-per-second; Gig-E 200, Library Inc., Tokyo, Japan) was located above the chamber (93 cm for pigeons, and 150 cm for crows). Using these experimental settings, we could video-record the pecking movements of the birds from a horizontal view for the two species for comparison.

**Figure 1.**
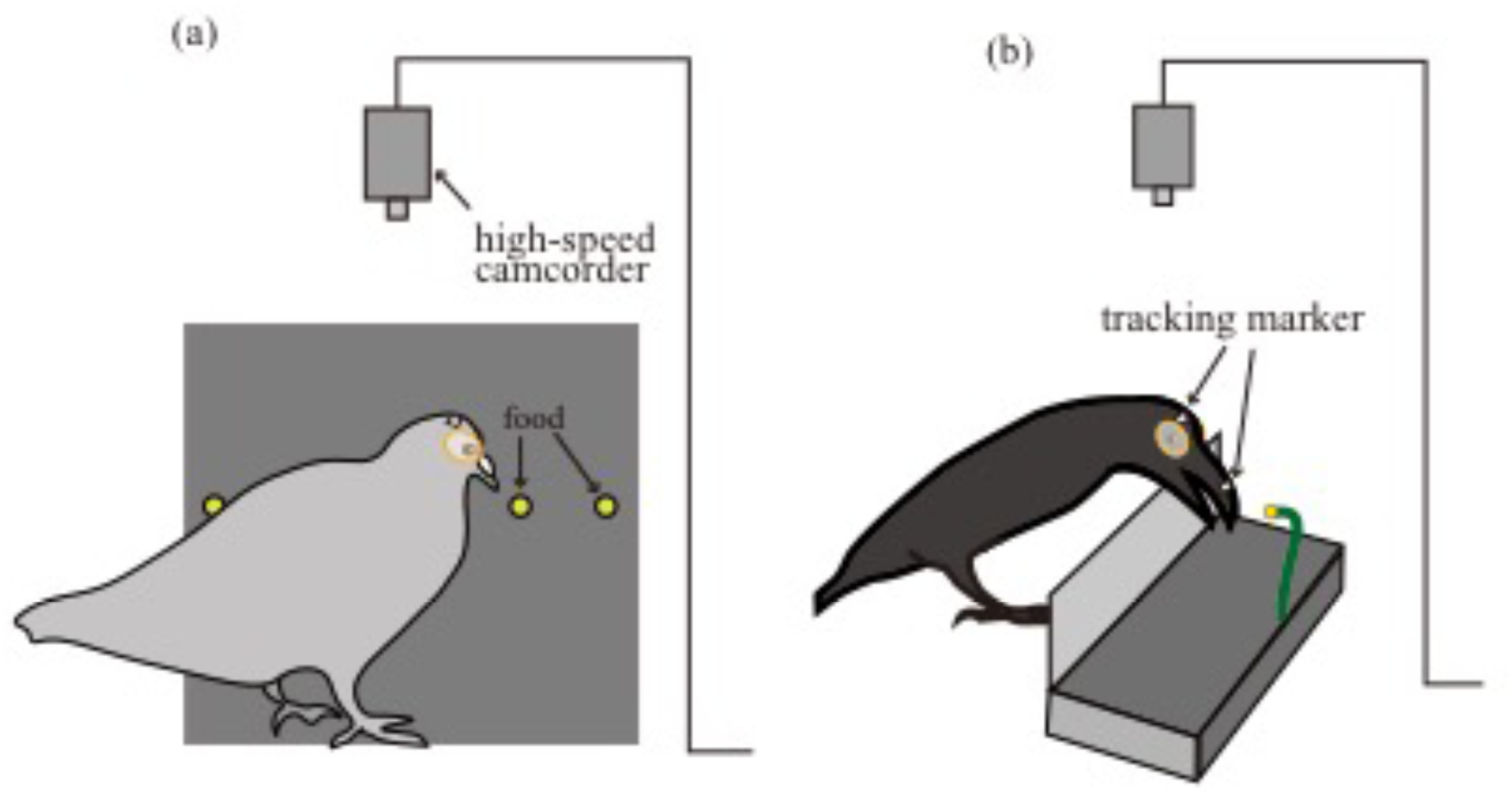
Schematic illustration of the experimental setting. Pecking movements were video-recorded with a high-speed camcorder mounted above the chamber to track horizontal coordinates of tracking markers (square-cut white pieces of tape, depicted in the figure). (a) For pigeons, five food items (corns) were attached to one of the walls of the chamber with adhesive tape. (b) For crows, the experimental setup was slightly different from that of pigeons. A crow pecked at a single food (a piece of cheese), which was attached to a wire on a platform.

**Figure 2.**
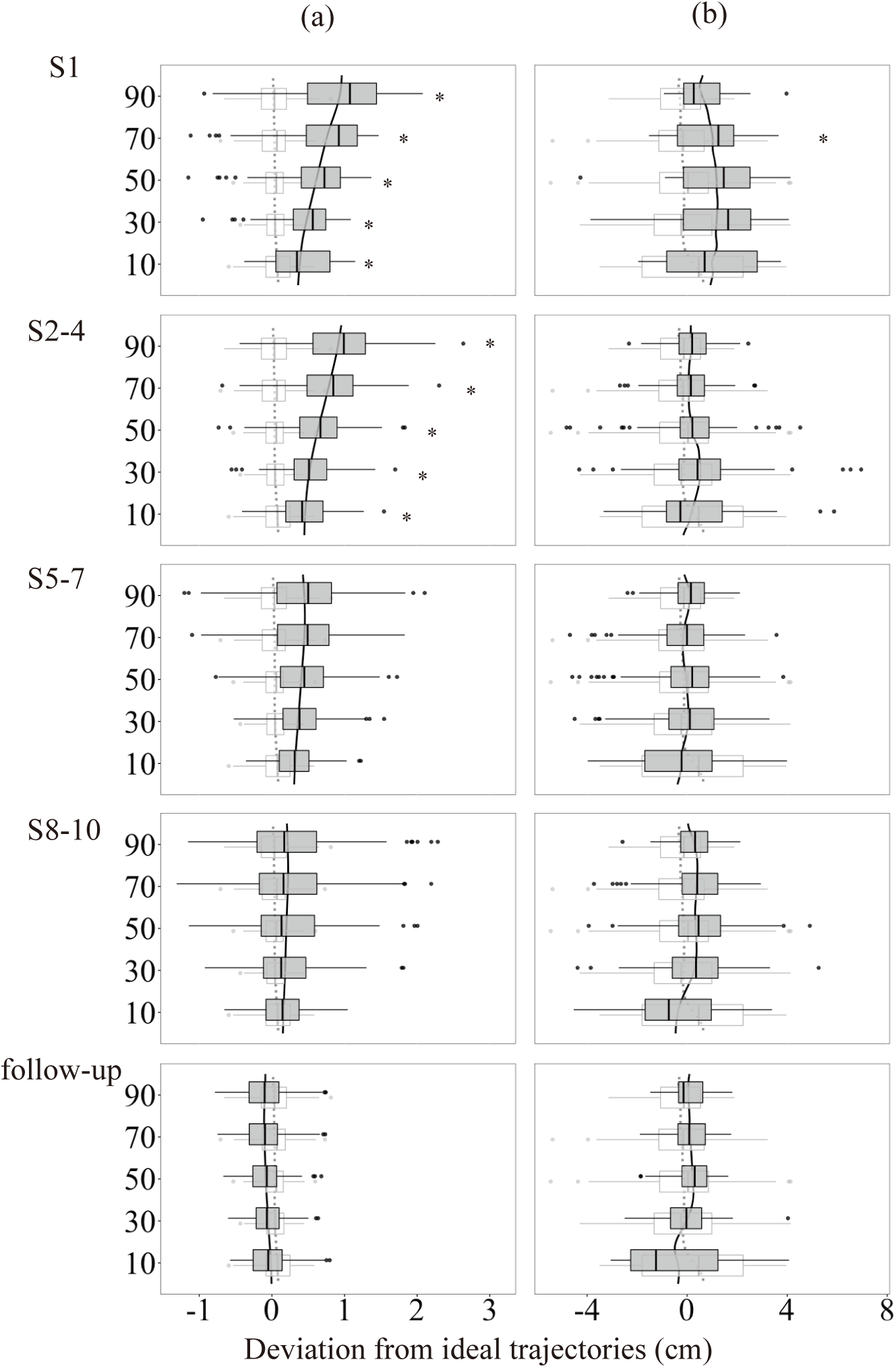
Lateral deviation from the ideal trajectories at each time point and phase. Positive values represent rightward deviation, and negative values represent leftward deviation. The translucent white boxes represent control phase. Asterisks indicate significant differences of deviation compared to the control phase. (a) data from pigeons. (b) data from crows.

### Procedures

For both pigeons and crows, the experiments consisted of the following three phases; *control* (normal eyes) phase (1 - 3 sessions), *prism* phase (10 sessions), and *follow-up* phase (1 session) phases. Each daily session consisted of 10 trials for pigeons and 20 trials for crows, which allowed us to record a maximum of 50 pecks from pigeons and 20 pecks from crows in each session. For all experimental phases, the session was terminated when the subjects consumed all foods or did not show responses for five minutes.

Before the control phase began, the subjects were briefly handled to attach the frames of prism glasses to their eyes with medical glue. The frames of glasses were made from dental resin, which had ellipse-like shapes (1.7 and 1.5 cm major and minor axis for pigeons; 2.0 and 1.7 cm for crows). The weights of glasses were approximately 2.5 g for pigeons and 3.5 g for crows. The procedure was performed under anaesthesia induced by inhalation of 3% isoflurane (Mylan Inc., Canonsburg, PA, USA). The control phase started at least after 24 hours after birds awoke from the anaesthesia, to allow the birds to recover from the procedure.

In the control phase, the subjects pecked at the foods without having the prisms attached to their glass frames. The control phase was performed for 1 - 3 sessions, until 50 instances of pecking were recorded for both pigeons and crows. The prism phase started the day after the control phase was completed. The filmy-thin prism (15-diopter; Fresnel prism, 3M company, MN, USA) was attached to the glass frames just before the prism session was begun. Anaesthesia was not used during prism attachment because the procedure was not considered to be painful, and took less than five minutes to complete. The prism was attached so as to shift the perceived position of foods 9.4˚ to the right, and remained attached after the daily experimental sessions. Thus, the subjects were involved in additional feeding and other daily activities in their cages whilst wearing prism glasses. Ten prism sessions were performed for each bird. The follow-up control session was conducted one week after the last prism session. Total experimental periods were 20-21 days, which included a week-long recovery period between the prism phase and the follow-up phase.

### Statistical Analysis

A single pecking movement was extracted from video images. We considered a pecking instance as a sequence of movements from head movement initiation to grasping offset. For the kinematic analysis, horizontal x-y coordinates of tracking markers were extracted using tracking software (Move-tr/2D v. 7.0, Library Inc., Tokyo, Japan). Small pieces of square-cut white tape were attached to the head for pigeons, and to the head and middle of the bill for crows, as tracking markers. Nose knobs were also used as tracking marker for pigeons. Thus, two tracking points were obtained for both species. The pecking instances in which tracking points were framed-out by head rotations were removed from the analysis.

Extracted coordinates were smoothed and segmented to 101 time points, using the smooth spline function. Trajectory segmentation was needed to conduct the analysis described below. We calculated horizontal deviation from the ideal trajectory in each time step. The ideal trajectories were defined as straight lines from initial head coordinates to food coordinates. The deviation for each time step was calculated as the distance of the perpendicular line from the bill tracking point (i.e. nose knobs for pigeons, and markers on the bills for crows) onto the ideal line. This measurement of deviation decreases when subjects approach the food in a straight line, while it increases when subjects deviate their movement orientation from the straight line to the food.

To examine whether deviation changed between phases, we compared deviation of each phase using linear mixed models. The time-points of 10, 30, 50, 70, and 90% of the pecking trajectories were chosen for comparison. The linear mixed model included the phases, time point, and the interaction between these as explanatory variables, and individual bird as a random effect. If the interaction was significant according to the likelihood ratio test, the same model analysis was performed for each time-points. The differences between phases were compared using 95% confidence intervals of estimated parameters of the model.

To examine the dependence of movement deviation between a time point and a preceding/subsequent one, correlations of deviations between all pairs of 101 time points were calculated and represented in a time-point space (Figure 3). Specifically, correlation coefficients were calculated for each pairs of time points by using pooled data of all pecking instances across phases. If the feedforward mechanisms were solely in control, which is the traditional view of pecking, the initial deviation would strongly influence subsequent deviations because the movement orientation was determined at the initiation of pecking but not adjusted during pecking. Thus, we predicted that high correlation would be observed between each deviation. On the contrary, if the on-line control mechanism at work, deviation would be reduced during movement, because the movement orientation to the target would be adjusted based on the preceding movement during pecking. Thus, we predicted that, for crows, which are thought to have on-line motor control, the correlation between deviations of each time points would be low.

**Figure 3.**
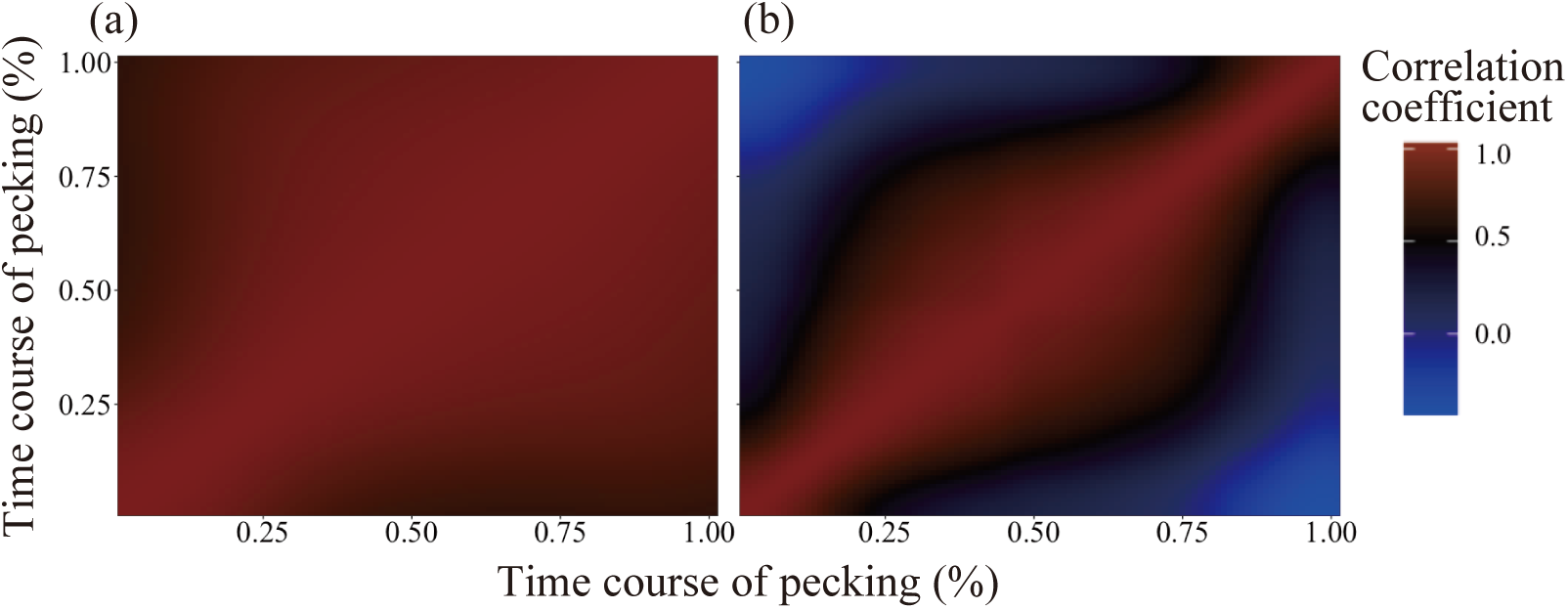
Heat-map representation of correlational analysis of deviation between each time-step. Colour reflects correlation coefficient. (a) data from pigeons. (b) data from crows.

All analyses were performed using R statistical software, version 3.4 (R Core Team, 2016). We used the ‘lme4’ package for the generalized linear model (Bates et al., 2015), and the ‘car’ package for the likelihood ratio test (Fox and Weisberg, 2011).

## RESULTS

### Pigeons

A total of 1,296 instances of pecking were recorded from the three pigeons. This included 150 instances in the control session, 80 in session (S) 1, 192 in S2 – 4, 303 in S5 – 7, 429 in S8 – 10, and 142 in the follow-up session.

The analysis of deviations revealed significant effects of the three phases (control, prism, and follow up, χ^2^= 86.94, *df* = 6, *p* < 0.001), and the interaction between time points (10, 30, 50, 70, and 90% of the pecking trajectories) and the phases (χ^2^ = 36.58, *df* = 5, *p* < 0.001). Subsequent analysis was performed on each time point, and revealed that the phase variable was significant across all time points (10%, χ^2^ = 267.96, *df* = 6, p < 0.001; 30%, χ^2^ =306.20, *df* = 6, p < 0.001; 50%, χ^2^ = 328.29, *df* = 6, p < 0.001; 70%, χ^2^ =373.26, *df* = 6, p < 0.001; 90%, χ^2^ =436.82, *df* = 6, p < 0.001). The phases that were significantly different from the control phase are depicted in Figure 2a.

Correlational analysis between each time step revealed strong correlations between each deviation (Figure 3a). The lowest correlation between 0% and 100% was 0.66. Correlation coefficients above 0.7 were observed in 96.6% pairs of time steps, suggesting that deviations of each time steps were strongly dependent on each other.

### Crows

A total of 624 instances of pecking were recorded from the three crows. This included 142 instances in the control, 32 in S1, 129 in S2 – 4, 123 in S5 – 7, 140 in S8 – 10, and 58 in the follow-up.

The analysis of deviations revealed significant effects of the three phases (χ^2^= 1616.44, *df* = 6, *p* < 0.001), and the interaction between time points (χ^2^ = 169.43, *df* = 5, *p* < 0.001). Subsequent analysis was performed on each time point, and revealed the phase variable was significant across all time points (10%, χ^2^ = 15.77, *df* = 6, p < 0.02; 30%, χ^2^ =33.66, *df* = 6, p < 0.001; 50%, χ^2^ = 27.98, *df* = 6, p < 0.001; 70%, χ^2^ =43.19, *df* = 6, p < 0.001; 90%, χ^2^ =64.90, *df* = 6, p < 0.001). The phases that were significantly different from the control phase are depicted in Figure 2b. The results suggest that the deviation increased only at the 70% time point of the first prism session, and was corrected for at the later 90% time point.

The correlational analysis in crows produced results that were very different from those of pigeons (Figure 3b). Crows showed high correlation only between adjacent time steps, while correlation decreased as time steps decreased. In particular, the correlation between the initial time steps (< 25%) and the final few steps (> 75%) was close to 0 or moderately negative (> −0.5). This suggests that the initial deviation scarcely influenced the final deviation in crows.

## DISCUSSION

The current study aimed to determine the controlling mechanisms of pecking in pigeons and crows by using prism-induced visual shifts. We found the contrasting results between crows and pigeons: crows rapidly adjusted the movement trajectory and correlations of the movement deviations were limited only between the adjacent time steps but very weak between the initial and the final steps. On the other hand, pigeons slowly adjusted the movement trajectory and the movement deviations were strongly correlated between the initial time step and the subsequent ones. These results suggest that pecking of pigeons and crows are under feedforward control and on-line feedback control, respectively. Our results provide the first evidence to suggest the on-line visuomotor control in pecking of birds.

Effects of prism at the levels of both kinematical and correlational analyses in pigeons support the traditional view of pecking mechanism (Goodale, 1982; LaMon and Zeigler, 1984; Matsui & Izawa, 2017), the feedforward control. On kinematic level, prism induced significant rightward deviation of movement trajectories until prism phases S2 – 4 (Figure 2a). The deviation gradually decreased in the subsequent S5 – 7 and S8 – 10 phases, indicating that pigeons learned to correct their movements to cope with prism-induced visual shifts. This gradual correction of movement trajectories through the phases is consistent with feedforward off-line control. Under feedforward control, the sensory signal for movement correction is obtained only after pecking completion, resulting in the learning with pecking-by-pecking updates like as we found in the kinematic analysis. The correlational analysis on deviations revealed that high correlation values were spread across the entire time step space (Figure 3a). This result reflects that deviation in the initial period (i.e. just after the pecking initiated) strongly influenced the deviation in the subsequent time points until the end. In other words, a movement trajectory and its deviation would be determined at the initiation of pecking but not be adjusted during movement. Thus, these results suggest the feedforward control of pecking in pigeons.

In crows, on the other hand, the on-line control is supported by both kinematic and correlational analyses. Even in the in the first session (S1) just after the prism attachment, rightward deviations of pecking trajectories occurred only at the 70% time point of the trajectory but converged around 0 at the 90% time point. In the subsequent phases, no clear deviation of trajectories was observed. Little effect by prism attachment was also confirmed by the correlational analysis. Positive correlations were found limitedly between adjacent time points but very weak among distant time points (Figure 3b), indicating that the initial deviation did not determine the final deviation. It is notable that little effect on pecking kinematics cannot be accounted by ineffective attachment of the prism because the success rate of pecking at the target food significantly dropped in both crows and pigeons (Figure S1a, b). Given that the prism actually distorted the perceptual position of target foods, the present results from crows suggest the on-line motor control to reach accurate end-points of pecking.

Past studies birds suggested that pecking is executed under feedforward control based on a motor command planned just before initiating pecking but under no on-line feedback control during movement (Goodale, 1982; Theunissen et al., 2017). Thus, the traditional view is that pecking in birds is ballistic and less flexible movement. This view is consistent with our present results from pigeons that movement trajectories were shifted in response to the prism and the adjustment of movement to the shifted vison occurred slowly with pecking-by-pecking updates. Poor adjustment of pecking to the prism-shifted vision was also found in a series of past studies reporting that chicks of domestic chickens failed to adjust pecking trajectories during up to 16-day sessions with attachment of prisms (Rossi, 1968; 1969; 1971). These findings together with our present one are consistent with the hypothesis of the off-line feedforward control in pecking.

On the other hand, the feedforward hypothesis cannot account for the present results of crows. Their rapid adjustment of movement trajectories and weak correlations of movement deviations during pecking are rather in accordance with the hypothesis of on-line feedback control. Although we did not determine the visual information and the time point/range critical for on-line feedback control, recent studies in carnivorous/omnivorous generalist birds, such as crows, suggest the active role of vision in control pecking or neck-reaching-based foraging behavior. New Caledonian crows, a tool-using bird in the wild, have been found to have extraordinarily large frontal-visual fields (Troscianko et al., 2012) and also the dominant side of the eyes (i.e. laterality) for tool use (Martinho et al., 2014). Large-billed crows, non-too-using species in the wild, those which were trained to use a hook tool to retrieve a food on a table, showed the deterioration of tool movement trajectories by blocking visual information of the table (i.e. tool tip and food) with an opaque cover (Kanai et al., 2014). Our present results of crows provide the first behavioural data to strongly support the on-line feedback control in pecking of birds, beyond the traditional hypothesis of avian pecking mechanisms, and provide the novel research direction of visuomotor mechanisms underlying the dexterous foraging skills, such as tool use in corvids, in carnivorous/omnivorous generalist avian species.

## ACKNOWLEDGMENTS

We thank the staff of the Biopsychology lab, Keio University for caring for the animals.

